# A Noninvasive Skin Biopsy of Free Nerve Endings via Realtime Third-Harmonic Microscopy

**DOI:** 10.1101/2022.12.14.520369

**Authors:** Pei-Jhe Wu, Hsiao-Chieh Tseng, Chi-Chao Chao, Yi-Hua Liao, Chen-Tung Yen, Wen-Ying Lin, Sung-Tsang Hsieh, Wei-Zen Sun, Chi-Kuang Sun

## Abstract

Skin biopsy was the only method to provide free-intraepidermal-nerve-endings (FINEs) structural information for the differential diagnosis of small fiber neuropathy (SFN). Its invasive nature was particularly unfavorable for patients with diabetic coagulation abnormalities thus there is an unmet clinical need for a non-invasive FINEs imaging tool. Here we show a tightly-focused epi-Third-harmonic-generation microscope (TFETM) for unmyelinated FINEs imaging. Its label-free capability was confirmed by PGP9.5 immunohistochemistry staining and a longitudinal spared nerve injury model study. Moreover, through proposing a dot-connecting algorithm, we established the operational protocol to count three-dimensionally the intraepidermal nerve fibers (IENF) and define the quantitative IENF index. Our clinical trial showed that the label-free IENF index can differentially identify SFN (*P*-value=0.0102) and was well correlated with IENF density of skin biopsy (Pearson’s correlation, R-value= 0.98) in the DPN group. Our study suggested that the unstained dot-connecting third-harmonic microscopy imaging can noninvasively provide FINEs structure information assisting diagnosing SFN.

## Introduction

Peripheral Neuropathy (PN) is the most common type of neuropathy, with hundreds of million population affected [1]. Diabetes is the primary cause of PN, accounting for 30%–40% of all PN cases [2-4]. Approximately 60% of all diabetic patients subsequently develop diabetic peripheral neuropathy (DPN) during their lifetime, thus prone to foot ulcers and even lower extremity amputations [5-9]. To avoid those DPN complications, the position statement of the American Diabetes Association recommends all individuals with diabetes should be annually screened because of the high prevalence of DPN [10,11]. Distal symmetric polyneuropathy is the most common type of DPN, and presents as symmetric numbness, paresthesia, and dysesthesias in the lower extremities, accounting for 75% of all DPN cases [12]. Small fiber neuropathy (SFN) represents an early stage of DPN, suggesting that terminal small nerve fibers of lower distal extremities are the early targets of DPN [13-15], thus our target for screening.

Diagnosis of SFN is currently challenging because physical examinations and painful nerve conduction studies primarily detect large sensory nerves and are ineffective for the functional assessments of small sensory nerves [16,17]. Quantitative sensory testing (QST) was commonly used for the functional assessments of sensory nerve fibers in SFN patients; however, it is psychophysically subjective and highly dependent on the experience of the experimenter and the testing environment [18-20]. Therefore, QST is more suitable for studying population changes over time than as a diagnostic tool for individual patients [21,22]. The assessment of intraepidermal nerve fiber (IENF) density has been widely deemed as the gold standard for diagnosing SFN. IENFs pathology with skin biopsy from the distal leg is routinely performed by immunohistochemistry using various neuronal proteins, particularly protein gene product 9.5 (PGP 9.5) [23,24]. Skin biopsy visualizes the number and morphology of IENFs, being supported by the European Federation of Neurological Societies [25-28], and the quantitative analysis of IENF density distinguishes SFN patients from normal subjects. Nevertheless, invasive skin biopsies, which could leave wounds on patients’ skin [29], are particularly unfavorable for patients with diabetic coagulation abnormalities. It was also difficult to standardize IHC staining, resulting in variability between laboratories [30]. Thus, label-free slide-free noninvasive imaging techniques for observing free intraepidermal nerve endings (FINEs) in the skin with its native contrast are essential and crucial to reducing harm to patients and potential medical risks.

Small sensory nerve fibers consist of thinly myelinated Aδ fibers, which transmit sensations of coldness and intense pain, and unmyelinated C fibers, which transmit sensations of warmness, hotness, and gradual pain [31,32]. Sensory nerve terminals are localized in the most superficial layer of the skin. [33] Morphologically, the axon diameter of unmyelinated FINEs is not fixed and ranges from 0.2 to 1.5 µm, based on electron micrograms. [34-36]. The sudden increase in axon diameter was found to be contributed by synapses rich in mitochondrial vesicles, glycogen granules, and other organelles, resulting in the characteristic varicosity appearance [37-39], thus providing the phase matching condition for tightly focused third-harmonic generation (THG). THG microscopy, as a noninvasive, label-free slide-free nonlinear imaging technique, is a powerful tool for the visualization of various cells and tissue structures and has been widely applied in neuroscience [40-45]. Due to the high sensitivity to lipid at a wavelength around 1180nm [46], 1180nm-based label-free THG microscopy has been applied to imaging the myelinated fibers in mouse brain tissues *ex vivo* and *in vivo* [47,48], and the THG signal was verified to be derived from the myelin sheath of Schwann cells [49,50]. Clinically, Florent Aptel et al. applied *ex vivo* THG microscopy to intact human ocular to visualize myelinated nerve fibers in the sub-basal layer of corneas [51]. To the best of our knowledge, all previous studies indicated the THG capability of imaging the myelin sheath of nerve fibers, with no demonstration and studies on *in vivo* clinical label-free imaging of unmyelinated FINEs either with THG microscopy or other imaging modalities.

In the present study, we propose and demonstrate a label-free method to noninvasively visualize unmyelinated FINEs through the tightly-focused THG microscopical imaging under 1260 nm broadband femtosecond excitation and develop a dot-connecting algorithm to realize *in vivo* label-free clinical structure imaging of FINEs. In a study using the nerve fiber staining agent PGP 9.5, *ex vivo* tightly-focused epi-THG microscopy (TFETM) was first confirmed to be capable to provide the label-free contrast of unmyelinated nerve fibers in the epidermis of human skin tissues of lower distal extremities. *In vivo* TEETM imaging of mice toe tips not only confirmed its imaging capability but also verified again that the signal is derived from the unmyelinated FINEs through a longitudinal study of a Spared Nerve Injury (SNI) small animal model. To enable the quantitative structure imaging capability of unmyelinated FINEs in human skin of lower distal extremities, we further developed an operational definition analysis based on a dot-connecting algorithm and proposed the intraepidermal nerve fiber (IENF) index. Our clinical study not only verified that the *in vivo* dot-connecting third-harmonic microscopical imaging obtained IENF index can differentiate between patients with DPN and control subjects, but also with high correlation with IENF density of skin biopsy in the DPN group. This unprecedented noninvasive methodology would fulfill the unmet clinical need for a label-free noninvasive quantitative FINEs imaging tool not only to assist the screening and differential diagnosis of DPN but also could for surgical evaluation and efficacy assessment of radiculopathy treatment and therapy in the future.

## Results

### Skin innervation of lower distal extremities tissue

The human skin tissue of lower distal extremities was richly innervated by nerve fibers immunoreactive for protein gene product 9.5 (PGP9.5) (Fig. 1b). Free intraepidermal nerve ending (FINEs) immunoreactive for PGP9.5 penetrated through the basement membrane after arising from the subepidermal nerve plexuses and had a typical varicose appearance (Fig. 1c). The unmyelinated FINEs elongated into the epidermis with multiple orientations and the morphology of FINEs is with a dot-like or string-like pattern which consists of dense varicosities (Fig. 1(d-e)). Moreover, the internal distance of each varicose was not fixed. The internal distance was 0 to 10.4 µm according to our staining results (Fig. 1(f-g)).

**Fig. 1.**
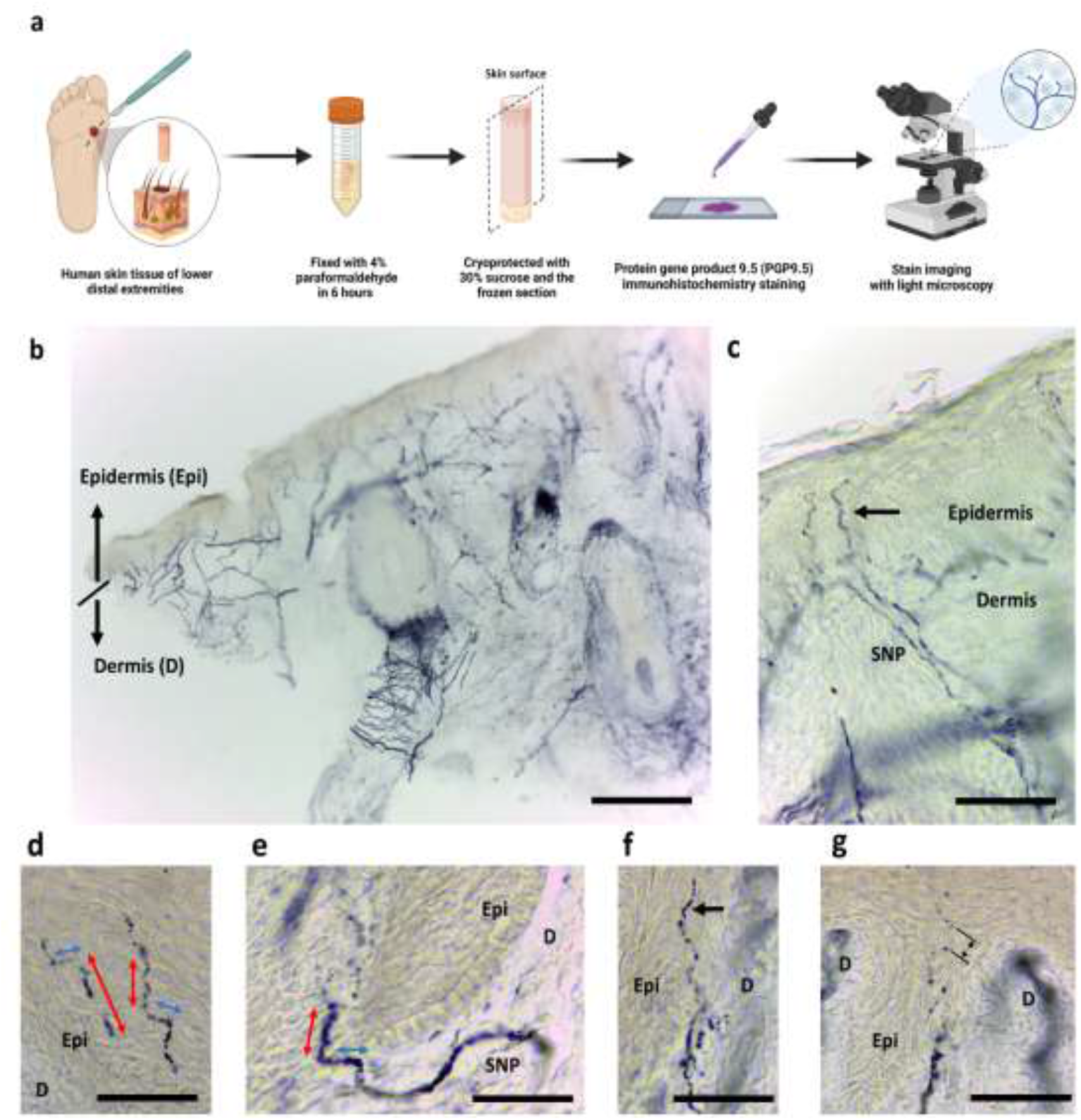
Skin innervation of lower distal extremities. (a) Human skin tissues were immunostained with protein gene product 9.5 (PGP 9.5). (b) PGP 9.5 (+) nerves are in the epidermis (Epi), and in the dermis(D). (c) Typical epidermal nerves arise from the subepidermal nerve plexus (SNP) and have a varicose appearance (arrow). (d-e) The Free intraepidermal nerve endings (FINEs) are with dot-like and string-like patterns at the epidermis with multiple extending orientations (red and blue arrows with orientation vertical and horizontal to the skin surface, respectively). (f-g) The range of distance between nearest neighbouring varicose bulges was 0 to 10.4 μm. Scale Bar =1 mm (b), 50 μm (c-g).

### Skin degeneration with post-fixed IHC staining

To avoid the influence of the fixation solution (4% paraformaldehyde) and the IHC staining on the epi-THG signal for histology validation, the post-fixed method was necessary. The fresh human skin section was cryoprotected with 30% sucrose and the frozen section was then conducted. The total duration required was 1 hour. We performed the post-fixed IHC staining 15, 30, and 60 minutes (resulting in accumulated delay for post-fixed staining: 1.25, 1.5, and 2 hours, respectively) after completing the TFETM imaging (Fig. 2a). Through the post-fixed IHC staining study, we learned that the FINEs would degenerate completely after 1.5 hours of delayed post-fixed staining (Fig. 2c). After 2 hours delay, the individual nerve fibers in the dermal nerve trunks either disappeared or the immunoreactivity became fragmented, indicating complete degeneration of nerve fibers (Fig. 2d). These results indicated that the *ex vivo* TFETM imaging for histology validation should be completed within 15 minutes right after the frozen section, and degeneration of FINEs should be expected (Fig. 2(b-d)).

**Fig. 2.**
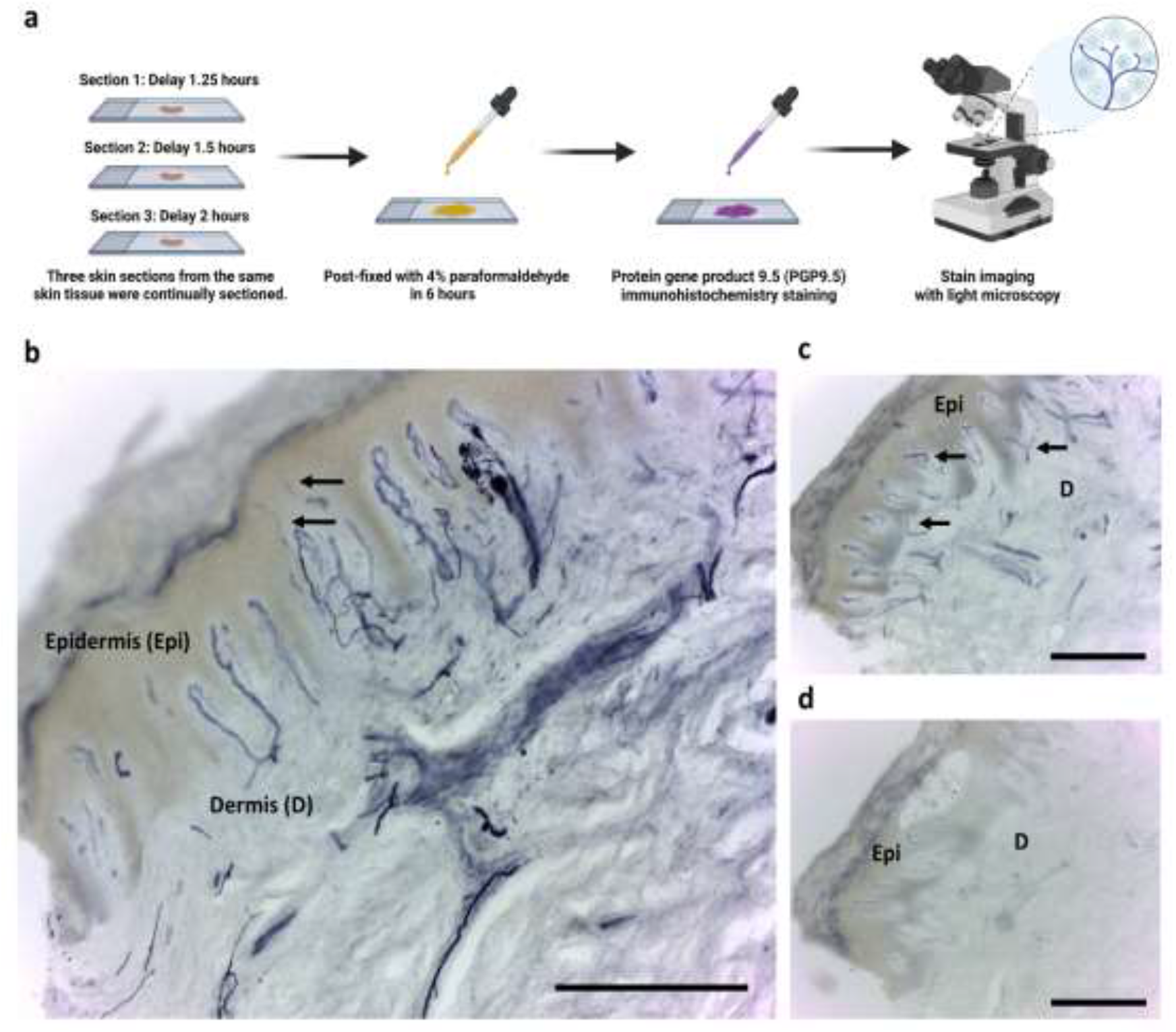
Skin degeneration of post-fixed IHC staining. (a) The post-fixed immunohistochemistry (IHC) staining in accumulated delay for post-fixed staining: 1.25, 1.5, and 2 hours, respectively. (b) The representative staining image of PGP 9.5-positive FINEs in the epidermis (Epi) (arrows) and dermis (D) after 1.25 hours of delayed post-fixed staining. (c) After 1.5 hours delay, only the nerve fibers in the dermis remained observed (arrows) and the immunoreactivity become fragmented in the epidermis. (d) After 2 hours delay, the individual nerve fibers in the epidermis and dermal nerve trunks disappeared, indicating complete degeneration of nerve fibers. Scale Bar =2 mm.

### *Ex vivo* tightly-focused epi-THG microscopy (TFETM) imaging of human skin tissue

For each unstained human skin section of lower distal extremities, the 3D stacks of *ex vivo* TFETM images which were composed of 2D *en face* images with different imaging depths at the epidermal-dermal junction (EDJ) were obtained (Fig. 3(e-m)). The label-free epi-THG signal showed many dot-like or pearl-like patterns (Fig. 3(l)), while stronger THG intensity and imaging contrast can be observed in structures with a string-like appearance consisted of closely packed dot-like signals in the epidermis. The epi-second-harmonic-generation (SHG) signal was generated specifically by collagen fibers in the dermis and effectively supported identifying the EDJ in the human skin section. Fig. 3(n) is the 2D overlapped image from Fig. 3(e-m), wherein epi-THG signals ascended perpendicularly from the dermis to the epidermis and had branches in the EDJ of the skin. With the 3D reconstruction image, the THG signal with a continuous linear pattern terminated in the upper portion of the granular layer of the skin was observed in Fig. 3(o-q). We define the optical propagation direction of the laser source as the z-axis. The TFETM not only can visualize the string-like or dot-like epi-THG signal with an orientation vertical to the optical axis but also the similar epi-THG signal with a structure orientation parallel to optical axis (Fig. 3(r-t)). To confirm the origin of the dot-like, pearl-like, and string-like THG signals, a histology validation with PGP 9.5 IHC staining was needed (next section).

**Fig. 3.**
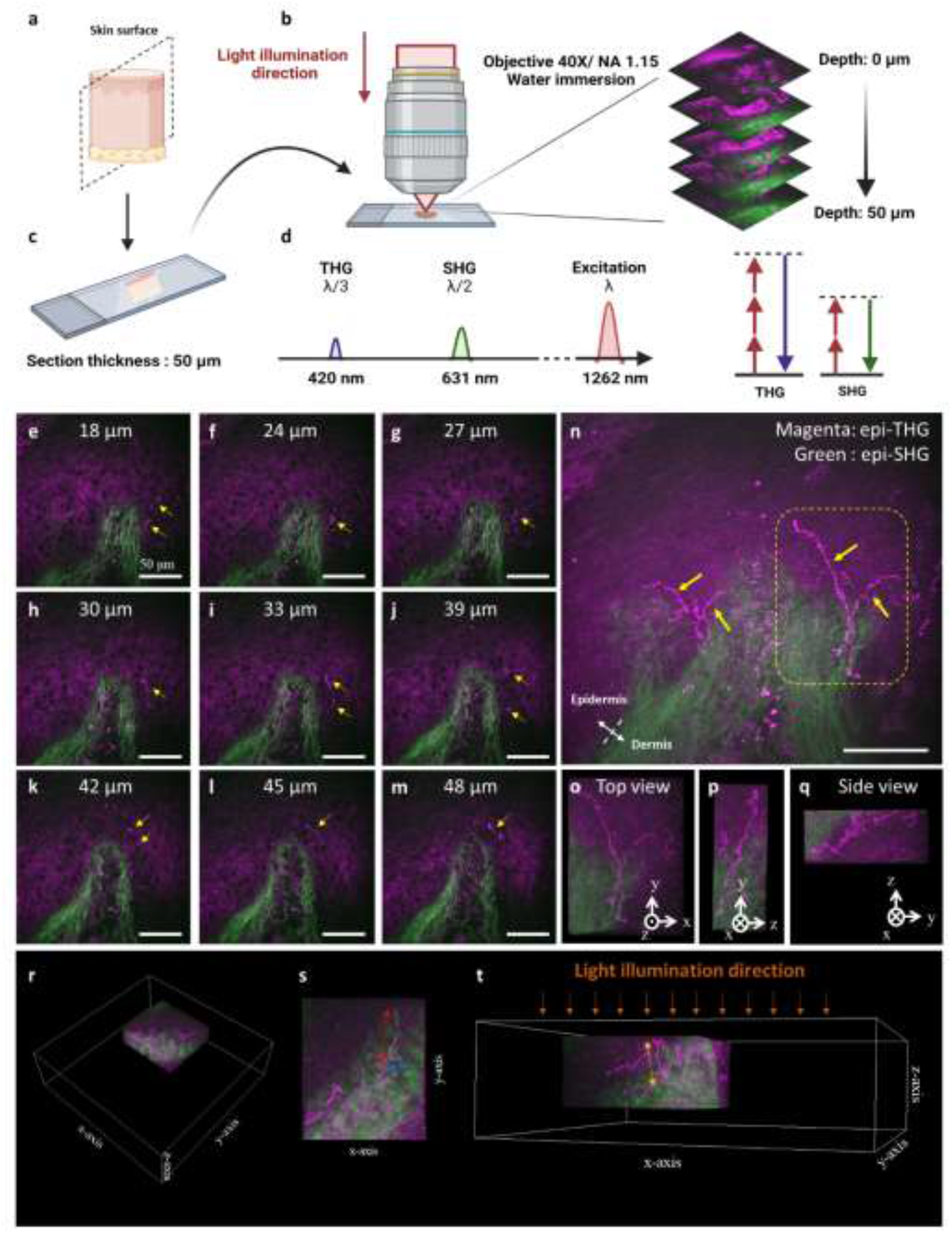
*Ex vivo* TFETM image of human skin tissue of lower distal extremities. (a) The human-distal region skin tissue was sectioned longitudinally. Dashed plane is the frozen section plane. (b) To achieve a tightly focused laser beam, a 1.15 NA objective was used to focus the laser beam onto the skin section. (c) The thickness of each skin section was 50 μm. (d) Spectral representation (left) of the signals created by excitation at λ=1262 nm. Third harmonic generation (THG) (resp. Second harmonic generation (SHG)) is generated at a third (resp. half) of the excitation wavelength. The corresponding simplified Jablonski diagrams (right). Dashed lines: virtual states. From (e) to (m), the 2D *en face* images with different imaging depths at the epidermal dermal junction (EDJ), and the epi-THG signal showed many dot-like or string-like patterns (yellow arrows) in the epidermis. (n) The 2D overlapped image from (e) to (m). The label free epi-THG signals ascended perpendicularly from the dermis to the epidermis and had branches in the EDJ of the skin. (o-q) The 3D reconstruction images showing the label free epi-THG signals with a continuous linear pattern in the epidermis of the skin. (r-t) The epi-THG signals with an orientation perpendicular (red and blue) or parallel (yellow) to the optical axis were observed. Scale bar = 50 μm.

### *Ex vivo* histology validation of human skin sections in lower distal extremities

Immediately after the *ex vivo* TFETM imaging, post-fixed IHC staining of the human skin sections with PGP 9.5 were conducted (Fig. 4a). Fig. 4 shows the representative results of the histological validation study performed on three individuals’ human skin tissues from lower distal extremities. Fig. 4(b-d) and Fig. 4(e-g) show the corresponding similar morphology and position of unmyelinated FINEs acquired by using the label-free TFETM imaging and by using immunolabeling imaging with PGP9.5 IHC staining from the same human skin sections. Fig. 4(h, j) show the zoom-in images. With higher molecule numbers of PGP9.5 IHC stains in FINEs structures with a larger volume, thus appearing with stronger blue color, the dot-like corresponding signals in both label-free TFETM and stained IHC images support the FINE origin of the dot-like epi-THG signals, but also suggest the epi-THG contrast on the varicosity structures of unmyelinated FINEs (Fig. 4(i, k)). Even though it is not background free, the strong epi-THG signals on the varicosity structures of unmyelinated FINEs provide a high contrast label-free image modality to provide morphological information of unmyelinated FINEs in the human epidermis with histological details.

**Fig. 4.**
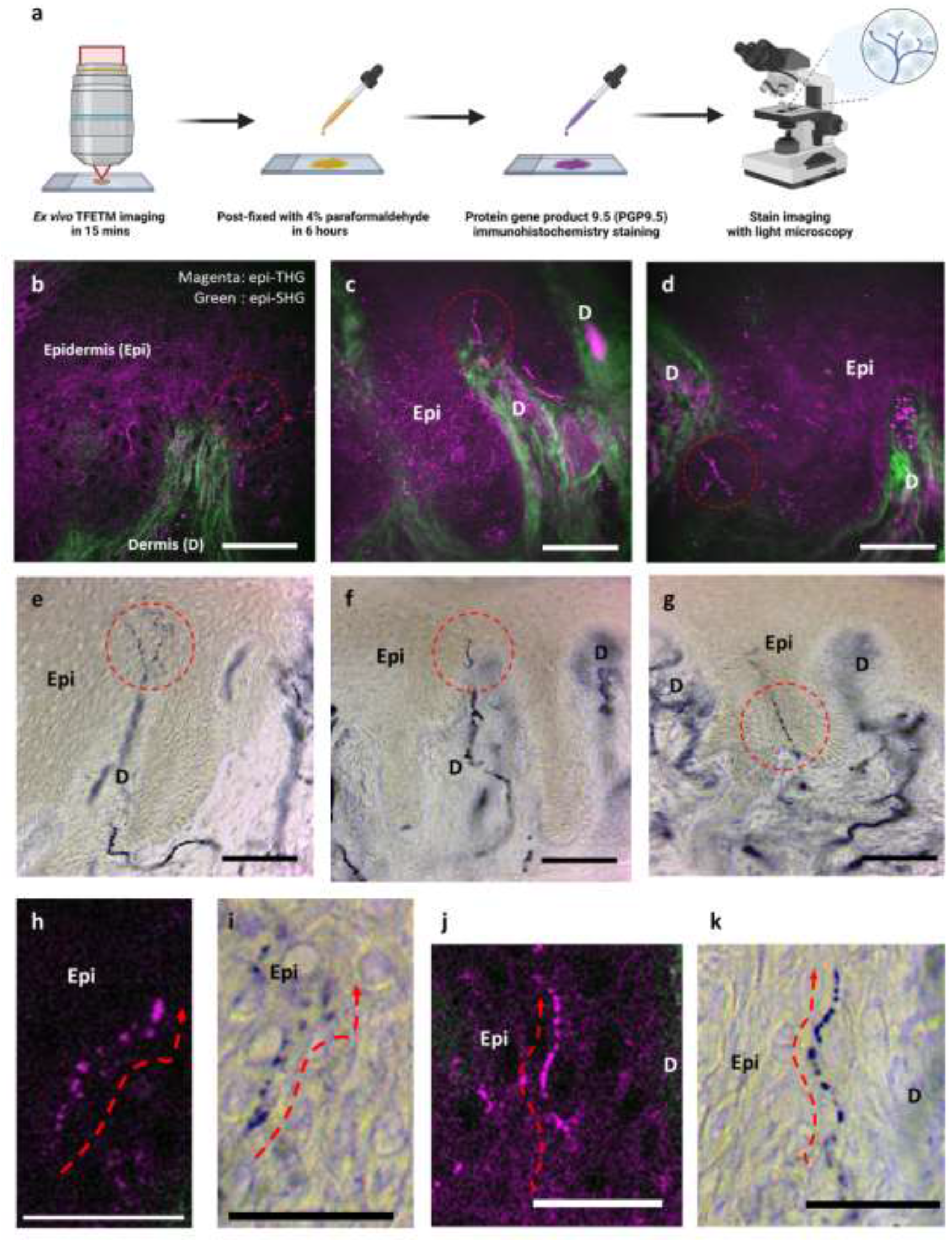
The representative results of the histological validation study performed on human skin tissues from lower distal extremities. (a) The post-fixed IHC staining of the human skin sections with PGP 9.5 were conducted immediately after the *ex vivo* TFETM imaging. (b–d) The unmyelinated FINEs with label free epi-THG signals (red circles) in the epidermis (Epi) of skin section by using the label-free TFETM imaging. (e-f) The corresponding similar morphology and position of unmyelinated FINEs acquired by using immunolabeling imaging with PGP9.5 IHC staining from the same human skin sections. (h, j) The label free epi-THG signal of the TFETM image appeared with a typical varicose structure of unmyelinated FINEs and was confirmed by PGP 9.5 IHC staining shown in (i, k). Scale bar: 50 μm. Epi: Epidermis. D: dermis.

### *In vivo* TFETM observation of nerve degeneration in wild-type mice following Spared Nerve Injury (SNI)

To further confirm the capability of *in vivo* TFETM imaging to visualize the FINEs, we used a partial SNI animal model with wild-type mice (C57BL/6J Narl mice) to observe the well-known damage process of FINEs in the mice toe tips (Fig. 5(b-c)). With three individual SNI mice, we performed *in vivo* TFETM imaging at a fixed acquisition position, which was the third toe tips of mice (Fig. 5(d)), at baseline, 24 and 48 hours after SNI surgery (Fig. 5(a)). At baseline, the dot-like or string-like epi-THG signals which was with similar morphology as *ex vivo* experiment was observed at the epidermis ((Fig. 5(e)), and also appeared simultaneously in three distinct mice (Fig. 5(f, i, l)). Within 24 hours of SNI surgery, the dot-like or string-like epi-THG signals, resembling varicosities of FINEs, can still be found in the epidermis of mice toe tips (Fig. 5(g, j, m). As expected from nerve innervation, after 48 hours, all the epidermal dot-like or string-like epi-THG signals disappeared in a stereotype fashion (Fig. 5(h, k, n)). These findings not only verified the *in vivo* TFETM imaging capability of FINEs, but also indicated the high TFETM contrast on the FINEs’ varicosities, thus with a dot-like THG appearance.

**Fig. 5.**
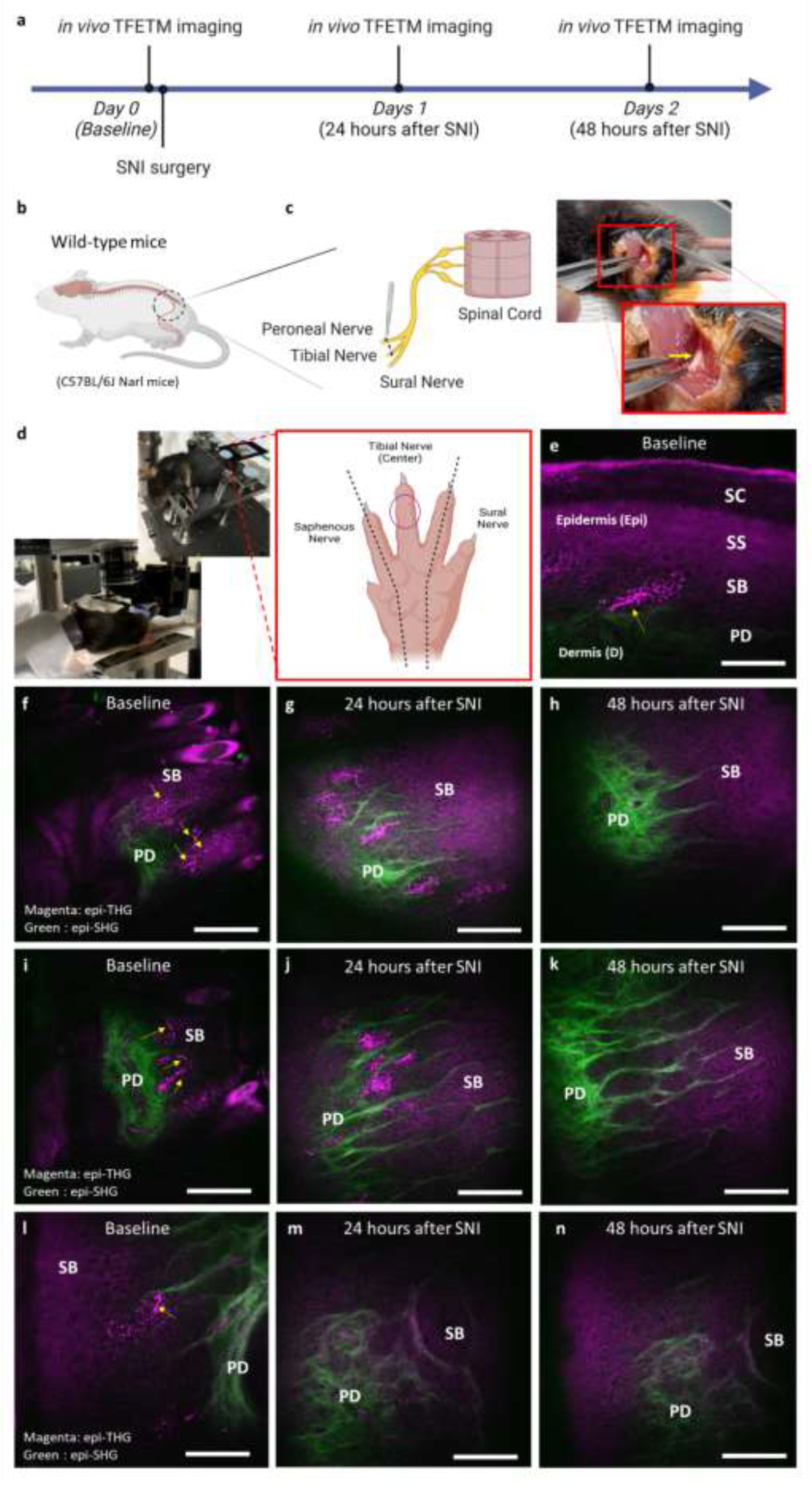
Longitudinal observation of nerve innervation and degeneration in mice toe tips before and after sciatic nerve injury (SNI) surgery with *in vivo* TFETM imaging. (a) The *in vivo* TFETM imaging at a fixed acquisition position was performed before SNI surgery (baseline), 24 and 48 hours after SNI surgery. (b)The wild-type mice (C57BL/6J Narl mice) to observe the well-known damage process of FINEs in the mice toe tips. (c) With the Spared Nerve Injury (SNI), the tibial nerve and common peroneal nerve are ligated and cut (yellow arrow). (d) The SNI procedure leads to complete denervation of the tibial (central) innervated area of the hind paw glabrous skin. The acquisition position for *in vivo* TFETM imaging was at third mice top tips (purple circle). (e) Before SNI surgery, the nerve fibers can be observed with epi-THG signals (yellow arrow) ascended from papillary dermis (PD) to stratum spinosum (SS) layers in the epidermis. (f, i, l) With three individual mice, the FINEs appeared to have a dot-like or string-like pattern (yellow arrow) in the epidermis of the third mice toe tip at baseline. After SNI surgery, the FINEs with epi-THG signal start to degenerate and completely disappeared after 24 hours (g, j, m) and after 48 hours (h, j, n). Scale bar: 50 μm. SB: stratum basale.

### *In vivo* TFETM imaging in human distal skin and IENF index

For clinical *in vivo* TFETM imaging, epidermal skin in the lower distal extremities is of the primary interest. Following an area selection study, the skin of instep arch was selected not only due to its relatively low melanin content in the basal layer, but also due to its shallower epidermal thickness (Methods, Imaging acquisition area selection for clinical *in vivo* TFETM imaging). Unmyelinated FINEs images, with string-like or dot-like structures either in the horizontal (Fig. 6(a–d)) or vertical direction (Fig. 6(e-f)), were successfully acquired by *in vivo* TFETM. With a dot-like appearance aligned/propagating in both horizontal and vertical directions and with the observation in all samples in all *ex vivo*/ *in vivo* human and animal studies, we conclude the sensitivity of TFETM to the varicose structure in FINEs (Fig. 6). For quantitative analysis, based on above conclusion and our PGP 9.5 IHC staining data, we further propose and developed the operational protocol base on a dot-connecting algorithm to identify the FINEs signal and provide the IENF index (Methods, Dot-connecting algorithm). For *in vivo* clinical study results, we conducted quantification analysis and compared the IENF index of different subgroups: the DPN groups (n=3) and the control subjects (n=11). The IENF index of *in vivo* TFETM image in DPN patients (mean ± SD: 19.7 ± 11.97 fibers/mm^2^) was significantly lower than control subjects (36.79 ± 7.79 fibers/mm^2^; *P* = 0.0102, paired *t*-test), as shown in Fig. 6(h). Moreover, the IENF index is significantly correlated with the IENF density of skin biopsy in three individual DPN patients (R-square=0.964, Pearson’s correlation: R-value=0.982, Fig. 6(i)).

**Fig. 6.**
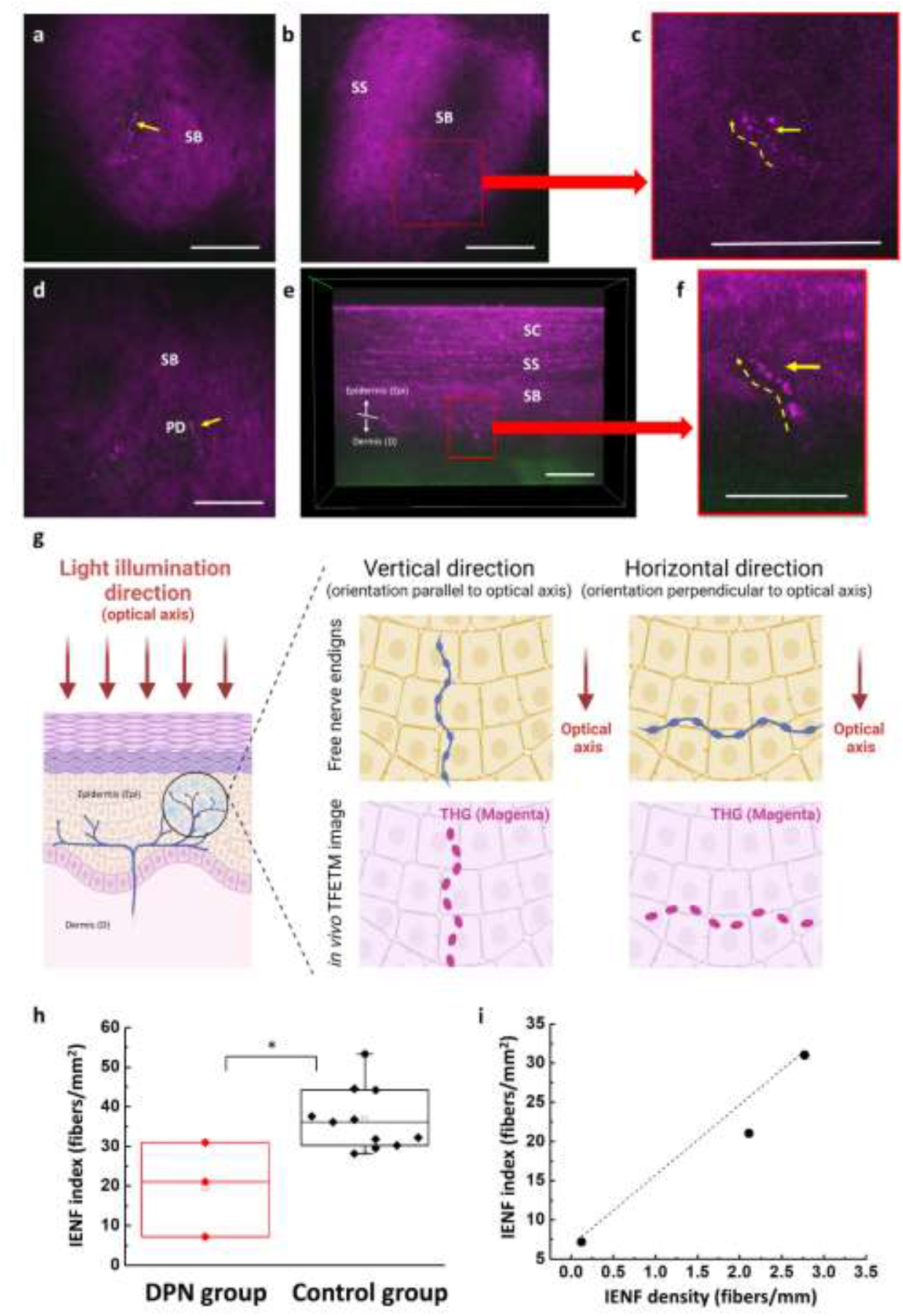
Representative *in vivo* TFETM image from the skin of instep arch in the lower distal extremities, and the quantitative analysis (IEFN index) in DPN and control subjects. (a, d) The label free epi-THG signal appeared and was with continuous dot-like or string-like pattern (yellow arrows) in the epidermis. (b, e) The unmyelinated FINEs images was with a typical varicose structure (red square) in the epidermis. (c, f) With the zoom-in images, those typical varicose structure of unmyelinated FINEs are with orientation horizontal and vertical to the skin surface, respectively. (g) The schematic diagram of *in vivo* imaging, the unmyelinated FINEs elongated into the epidermis with both horizontal and longitudinal directions to the optical axis can be visualized by TFETM. (h) In quantitative analysis with a dot-connecting algorithm, the IENF index of DPN patients with significantly lower compared with the control subject (\**P*<0.05, \*\**P*<0.01) and highly correlated (Pearson’s correlation R-value = 0.98) with IENF density of skin biopsy. Scale bar: 50 μm.

## Discussion

The neurites are periodically varicose, especially for FINEs. Our studies, including the study with the *ex vivo* study with the IHC staining, indicate the high sensitivity of TFETM toward the bulges, with sudden increase in axon diameter, of the varicose FINEs. PGP 9.5, a ubiquitin carboxy C-terminal hydrolase [51], is particularly enriched in small-diameter nerves and is thus with a higher total volume in the enlarged bulges compared to their surroundings. Our histopathological validation study comparing epi-THG and PGP9.5 IHC staining signals shows corresponding dot-like structures, thus supporting the contrast of THG as the bulges of the varicose FINEs. Morphologically, electron micrograms had reported the maximum bulge diameter of the varicose FINEs to be less than 1.5 µm [34-39], which is slightly smaller than the confocal parameter of focused laser beam after the objective of our designed TFETM so as to be able to provide the phase matching condition for bulge size and its provision on the spatial inhomogeneity required to generation strong THG out of its normal background. The THG phenomenon is a nonlinear coherent optical process induced by structures with specific properties. It’s well-known that the signal contrast is generated at material interfaces where there is a spatial distribution change in optical refractive index or third-order nonlinear susceptibility (χ3) within the confocal parameter. As THG is a third-order process, ultra-short laser pulses with high peak power densities at the optical focus are required to ensure sufficient signal generation. Theoretically, with a uniform-diameter fiber elongated toward the direction of the light propagation, the Guoy phase-shift effect will lead to the coherent cancellation of THG due to axial symmetry. Our observation of bulges indicates the origin of the tightly-focused THG as the varicosity of FINEs. The varicosity of FINEs with an enlarged appearance but with a size well-within the confocal parameter destroys the structural axial symmetry and results in optical inhomogeneity required to generate THG. The TFETM images of unmyelinated FINEs further indicated that the THG contrast on FINEs varicosity is independent of the nerve fiber orientation (Fig. 3(r-t) & Fig. 6) either along, perpendicular to, or with an angle with the optical axis.

With longitudinal tracking in the SNI animal model, the observed dot-like epi-THG signals at baseline disappeared completely 48 hours after nerve injury. This phenomenom is consistent with previous studies [53,54], which indicated the degeneration of FINEs 48 hours after nerve injury, thus confirmed again that dot-like signals in noninvasive TFETM are to *in vivo* reflect the contrast of the unmyelinated FINEs. In the *in vivo* clinical study, we demonstrate that the *in vivo* TFETM with a tightly focused NIR laser beam enables label-free imaging of unmyelinated FINEs. Furthermore, through the development of a dot-connecting protocol and algorithm, the IENF index of *in vivo* label-free TFETM was found to be able to differentiate between patients with DPN and control subjects non-invasively. Also, the IENF index was highly correlated with the IENF density of the skin biopsy in the studied group of DPN patients. It is noted that the epi-SHG modality, with a contrast of connective tissues that is the characteristic of the dermis layer, would assist the identification of dermal-epidermis junction. Our findings support our proposal on the development of the dot-connecting third harmonic microscopical imaging as a noninvasive label-free, quantitative, and objective FINEs imaging tool for the differential diagnosis of DPN and with high potential application for screening and assessment for radiculopathy treatment in the future.

### Langerhans cells and melanocyte dendrites in human skin

There are some similar morphological structures with nerve fibers that appeared in string or linear patterns in human skin including Langerhans cells and melanocyte dendrites. Our previous study confirmed that 1260 nm based epi-THG microscopy is not sensitive to dendrites of Langerhans cells [55]. Since THG is also sensitive to melanocyte dendrites, it is thus advised to choose skin areas with minimized melanocyte activities [56] and to use morphological characteristics to avoid the misdiagnosis of melanocyte dendrites as nerves. The melanocyte dendrites, an indication of hyperactivity of melanocytes, generally appeared on pigmented skin lesions, melanocyte tumors, and the skin after laser therapy rather than in normal skin [56].

### Corneal confocal microscopy (CCM) in neuropathic patients

The CCM has emerged as a *in vivo* ophthalmic imaging modality to evaluate small nerve fiber morphometry in Bowman’s layer, which is located below the basal layer in the cornea [57,58]. Several studies have shown that CCM has the capability to detect damaged nerve fibers in neuropathic patients [59,60], and that it demonstrated comparable diagnostic efficiency with skin biopsy [61]. However, CCM still reflects the contrast of myelinated axons. Furthermore, Ziegler et al. (2014) reported that although CCM and skin biopsy both detect nerve fiber loss in recently diagnosed type 2 diabetes, but largely in different patients, suggesting low correlation between CCM and IENF density of skin biopsy [62]. These results may indicate that small nerve fiber neuropathy does not develop simultaneously in different organs; thus, the results of nerve fiber damage in the cornea cannot directly reflect that in the skin of distal extremities. However, the most typical form of DPN is chronic, in which nerve fiber damage starts from the distal limbs and then spreads to the proximal limbs. This fact suggests that distal extremities skin is the earliest target of DPN. Our findings suggest the dot-connecting third harmonic microscopical imaging as a more reliable and suitable tool for label-free imaging of FINEs noninvasively in human skin, especially of lower distal extremities, for the differential diagnosis of DPN patients.

### Other noninvasive optical imaging techniques

Optical coherence tomography (OCT) can be used to obtain histopathological skin images with a high penetration capability. However, because OCT lacks spatial resolution, the sub-micron scale of neural information on FINEs cannot be acquired [63,64]. Photoacoustic microscopy (PAM) can achieve the ability to visualize myelinated mouse sciatic nerves *ex vivo*, and myelin can produce photoacoustic signals [65]. Reflection confocal microscopy (RCM) can provide information about myelinated nerves in deep brain regions of mice [66] or transgenic zebrafish [67] with strong contrast due to the high refractive index of lipid-rich myelin. Although PAM and RCM provide images with a higher resolution in human skin and seem attractive choices, there is no evidence to show that they can image unmyelinated nerves fiber, especially distal nerve endings C fibers.

In summary, FINEs are the most abundant sense organ to detect temperature, touch, and painful stimuli. Currently, pathological diagnosis based on surgically excised tissue followed by formalin-fixed and histochemical staining is the gold standard in clinical practice. While the affected skin and nerve is removed, longitudinal observation on the same targeted FINEs is not permitted. Here we show a noninvasive kin biopsy via TFETM for unmyelinated FINEs imaging. The proposed TFETM was shown to be able to delineate the 3D structure of unmyelinated FINEs that are confirmed by PGP 9.5 immunohistochemistry staining from the same tissue section. We then developed a dot-connecting algorithm to count the entire IENF number that are perfectly correlated with the quantitative IENF density by summation of all sections under light microscopy. Thus, we propose that TFETM is a paradigm-shift imaging to quantitative IENF density of traditional skin biopsy. Moreover, the real-time, non-invasive imaging technique will serve as a novel and powerful monitor to the regeneration and degeneration of FINEs throughout the clinical course of diabetic, chemotherapy-induced, postherpetic neuropathy.

## Methods

### Tightly-focused epi-THG microscopy (TFETM)

The TFETM system combined intravital epi-THG and epi-SHG and a femtosecond pulse laser with a central wavelength of 1262 nm, which was not only located within the transmission window of human skin but also avoids melanin absorption and skin scattering [68]. Therefore, the optical penetration depth can reach the epidermal-dermal junction even in the human-distal skin or glabrous skin. In the TFETM system, the 1262-nm Cr: forsterite laser, with a 105-MHz repetition rate and a 91-nm bandwidth, was used as the excitation source. Double-chirped mirrors were applied to control the optical dispersion. The laser beam was collimated and guided with a light guided (1.25 Series Laser Articulated Arms; LaserMech, Michigan, USA) into galvo-resonant mirrors (Laser Scanning Essentials Kit; Thorlabs, Newton, NJ, USA) to perform 2D scanning. To achieve a tightly focused laser beam, a 1.15 NA objective (40×; working distance: 250 μm; UApo N 340; Olympus, Tokyo, Japan) was used to focus the laser beam onto the skin. The backward SHG (∼630 nm) and THG (∼420 nm) signals were collected using the same objective and were received by two separate photomultiplier tubes (R928 for epi-SHG and R4220P for epi-THG; Hamamatsu Photonics, Hamamatsu City, Japan). Band-pass filters with different center wavelengths and bandwidths (FF02-617/73 for epi-SHG and FF01-417/60 for epi-THG, Semrock) were inserted before the Photomultiplier tubes (PMTs) to filter out the background noise to increase the signal-to-noise ratio. The objective was attached to a 3D step motor (TSDM40-15X, Sigma Koki, Japan) such that the 3D position of the objective could be adjusted by both manual and electrical controls. The imaging plane was moved to different depths by tuning the 3D stage along the optical axis. Following a similar pulse width measurement and dispersion control [69], the pulse width after the objective was estimated to be 28 fs. The epi-THG transverse and axial resolutions were approximately 0.4 and 1.1 μm, respectively [70].

### Imaging procedure

All the TFETM image stacks with a 14-bit grayscale were composed of a series of 2D *en face* images (Pixels size: 512 × 512, Field of view (FOV): 235 × 235 μm). The acquisition time was approximately 0.38 s per averaged image, after averaging five frames at a fixed depth with a 15-Hz frame rate. *Ex vivo* and *in vivo* TFETM images of depths from the surface was acquired every 0.6 and 1.8 µm along the optical axis, respectively.

### *Ex vivo* TFETM imaging in human skin sections of lower distal extremities

The study protocol was approved by the Research Ethics Committee of National Taiwan University Hospital (No. 200903064D), and informed consent was provided by all patients before study entry. The human-distal region skin tissue was cryoprotected with 30% sucrose and continually sectioned (thickness: 50 µm) using a cryostat (Leica CM 1950, Leica Biosystems, Wetzlar, Germany) for a total duration of 1 hour. For each skin section, the suspected areas with FINEs-like signals at the epidermis or epidermal-dermal junction were imaged and the acquisition position was recorded immediately.

### Immunocytochemistry (IHC) staining

After *ex vivo* TFETM imaging for a total duration of 15 minutes, human skin sections were immediately subjected to PGP9.5 IHC staining according to the procedure described in a previous study [71]. The skin sections were fixed with 4% paraformaldehyde for 6 hours, treated with 0.1 M phosphate buffer (PB; pH 7.4), and then stored for 12 hours. Furthermore, the skin sections were quenched with 1% H_2_O_2_, blocked with 5% normal goat serum, and incubated with rabbit antiserum to PGP 9.5 staining (UltraClone, UK, diluted 1:500 in 1% normal serum/Tris) at 4 °C for 16–18 h. After rinsing in Tris, and followed by incubation with the avidin-biotin complex (Vector, Burlingame, CA) for 40 min, the sections were incubated with biotinylated goat anti-rabbit immunoglobulin G at room temperature for 1 hour. The reaction product was visualized using chromogen SG (Vector). PGP 9.5-immunoreactive nerve fibers present in the epidermis of each section were counted at a magnification of 40× under a light microscope (Leica ICC50D, Leica, Germany).

### Spared Nerve Injury (SNI) small animal model

Wild-type C57BL/6J Narl mice were purchased from the National Laboratory Animal Center (Taipei, Taiwan). Mice aged 8–16 weeks and with an average weight of 25 g were used in this study. All mice were housed under a 12 hr light–dark cycle at a temperature of 22°C with food and water provided ad libitum. Both male and female mice were used in this study. All animal care and experimental procedures were approved by the Institutional Animal Care and Use Committee of National Taiwan University (approval number: NTU107-EL-00176). Mice were anesthetized using ketamine (50 mg/kg) and xylazine (15 mg/kg). The SNI surgery procedure was performed according to Decoster & Woolf (2000) and Burqini et al. (2006) [72,73]. Briefly, the fur on the left thigh was shaved. The skin and muscle near the knee were incised, and the left sciatic nerve was exposed. The common peroneal and tibial nerves, the two main branches of the sciatic nerve, were tightly ligated using a 6/0 silk suture, and a section of the two nerves was cut out using corneal scissors. The sural nerve was left intact. The muscle and the skin were then sutured using 6/0 silk. Lincomycin hydrochloride (30 mg/kg) was administered into the right gastrocnemius muscle to prevent infection. For the observation of epidermal innervation on the third toe using the TFETM, we recorded the acquisition positions at baseline, 24 hours, and 48 hours follow-up after SNI surgery.

### DPN Patients and control subjects

Three female DPN patients aged 62–70 (mean age: 67 ± 4.36) years were clinically diagnosed and were recruited. We also recruited eleven control subjects (three men and eight women) aged 24–62 (mean: 35.18 ± 13.36) years who were confirmed without diabetes and neuropathy based on glycated hemoglobin (HbA_1C_) blood test reports and neurological examinations. The study protocol was approved by the Research Ethics Committee of National Taiwan University Hospital (No. 201610037DINC) and the Taiwan Food and Drug Administration. Written informed consent was provided by all patients before the experiment. This trial was conducted according to the principles of the Declaration of Helsinki. The *in vivo* TFETM imaging was performed at the Molecular Imaging Center of National Taiwan University.

### Imaging acquisition area selection for clinical *in vivo* TFETM imaging

According to a previous study, the THG intensity is sensitive to melanin, while the melanin mass density (MMD) of the focal volume is directly related to the intensity of the THG signal in the basal layer [74]. In this study, although the THG signal was also found to be highly sensitive to the presence of FINEs, nerve fibers are mostly distributed among the basal keratinocytes, causing it to be easily shadowed by the surrounding background THG from melanin. For diabetic neuropathy patients, the most common symptom site is the distal extremities and with a length-dependent process, thus the acquisition area with a low melanin content on the distal leg was the optimal site for TFTEM imaging. Fig. 7(a-f) shows the TFETM images acquired at candidate areas on the terminal of the distal leg including the sole, heel, instep arch, bridge, and ankle (Fig. 7(g)). For positions 1, 2, 5, and 6, high melanin content in basal layers can be found to provide strong THG background. In contrast, positions 3 and 4 had little melanin in their basal layers. Our study concluded the instep arch to be the optimal choice. Moreover, the sole and heel of the foot caused serious light scattering due to the excessively thick stratum corneum and a much longer working distance objective will be required.

**Fig. 7.**
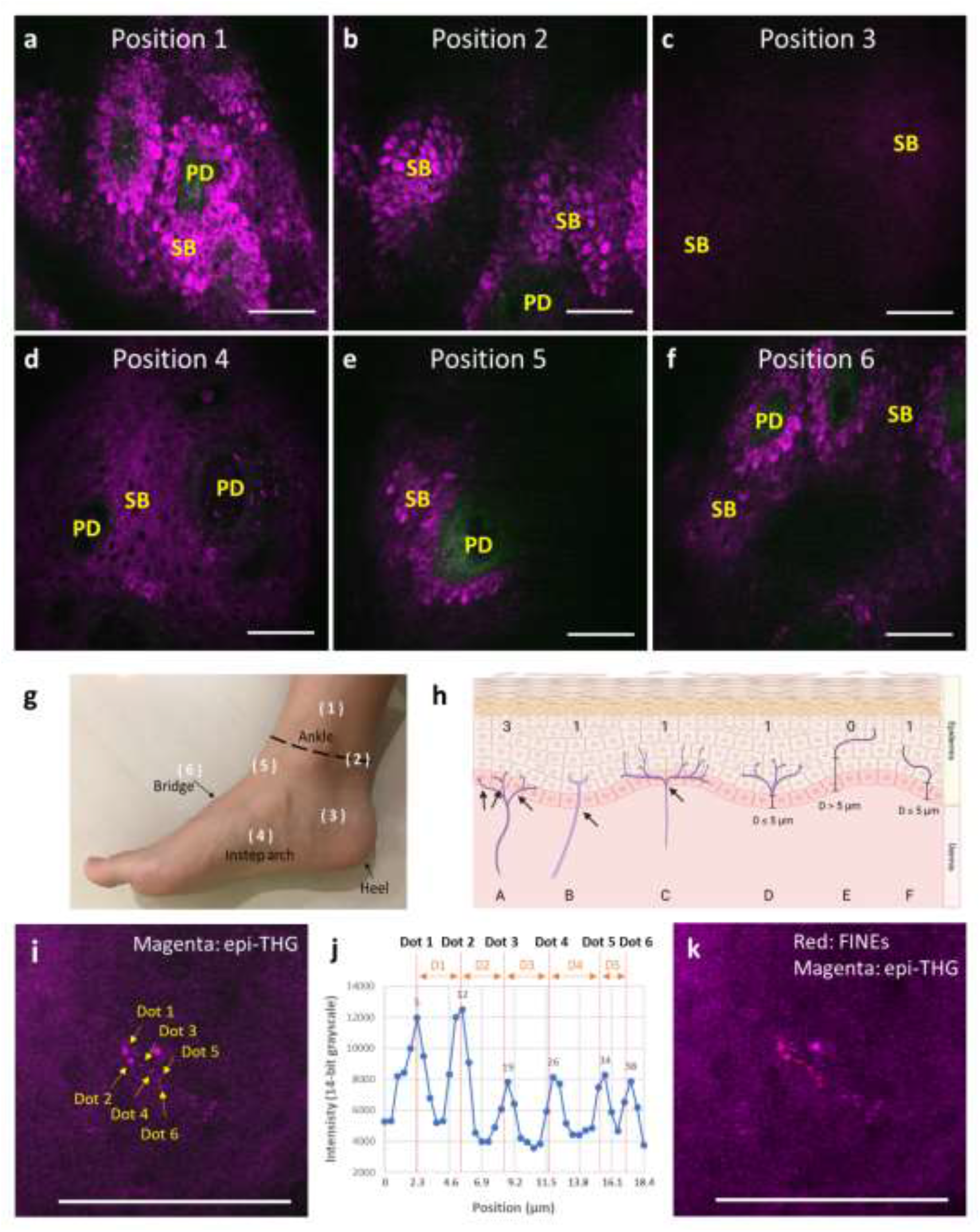
The *in vivo* TFETM images were acquired from different positions of the distal leg (a-f). (a, b, e, f) High melanin content in basal layers generates strong epi-THG background. (c, d) Much reduced melanin content in the basal layer can be found in positions 3 & 4, while the background epi-THG signal from position 3 was weaker than position 4. (g) The corresponding position of (a-f) on the foot. Position 1 was above the ankle; position 2 was on the ankle; position 3 was on the side of the heel; position 4 was at the instep arch; position 5 was between the ankle and the bridge; position 6 was on the bridge. (h) The diagram of skin innervation. FINEs are shown in magenta. (Case A) The FINEs with branching points in the epidermal-dermal junction (EDJ) or dermis were counted separately. (Cases B, C) The FINEs with a branching point in the epidermis was considered as one IENF. (E) The fragment, with a distance between the epidermal nerve fragment and the EDJ layer greater than 5 μm, was not counted. (Cases D, F) If the distance between the epidermal nerve and the EDJ layer was less than 5 μm, the IENF is counted. (i) A representative FINE signal showing at least 5 consecutive dot-like signals. (j) The internal distance between two successive dots-like signals should be within 10.4 μm. The internal distance of D1 to D5 was 3.21, 3.21, 3.21, 3.67, 1.83μm, respectively. (k) The same *in vivo* TFETM image as (i) was dot-connected (red line) as one IENF signal. Scale bar: 50 μm.

### *In vivo* TFETM of lower distal extremities

All the *in vivo* TFETM image stacks were acquired using the same imaging condition and at least twenty image stacks for each subject were acquired from the instep arch of lower distal extremities. The average laser power after the objective was 105 mW and the voltages of the photomultiplier tubes for the epi-SHG and epi-THG modalities were set at 700 and 800 V, respectively. Each image stack comprising 150 depth-dependent 2D sub-images with two channels, was obtained through the skin surface and upper dermis. We selected 1.8 μm as the z-step size for the optical section. To ensure a 250-μm-depth stack, 150 steps were acquired (under the 1.8-μm step size) with a total of 55 s. We also used an objective adaptor with a medical tape (3M™ Double Coated Medical Tape No. 9874) to fix and stabilize the relative position of the imaged skin and avoid motion artifacts. The allowed acquisition time for one patient was <40 min. Therefore, the total laser energy accumulated in the same area was <200 J. Photodamage evaluation and discussion under a similar clinical condition have been described elsewhere [75], while the same result was obtained in this present study. In the present study, we found no erythema, pigmentation, or blister formation on the examined skin.

### Dot-connecting algorithm

The operational protocol of the dot-connecting algorithm was established to identify the FINEs and count the number of IENFs within the TFETM imaging area. The details are summarized in the following statement and the flow chart shown in Fig. 8.

**Fig. 8.**
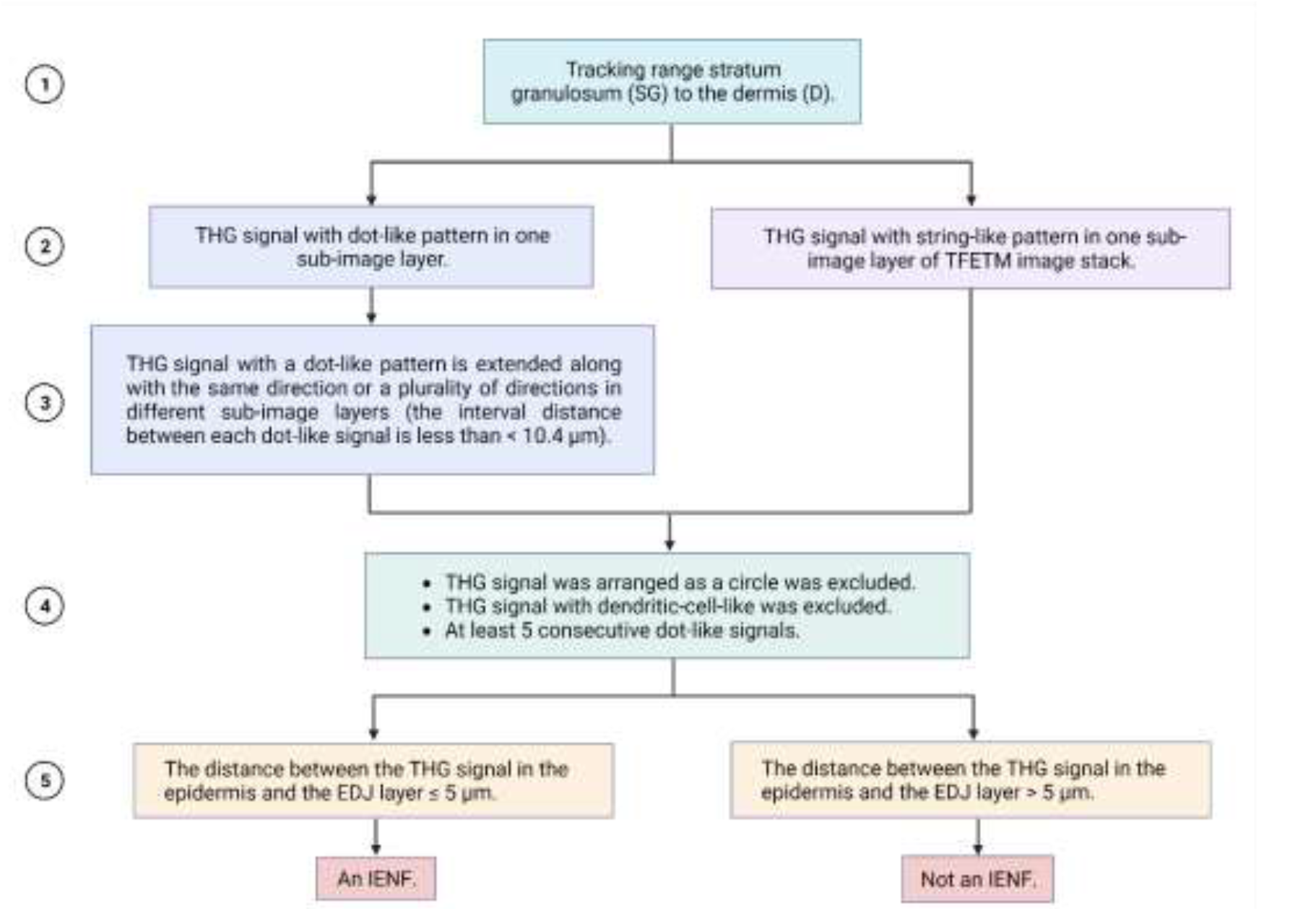
The flow chart of the operational protocol of dot-connecting algorithm.

#### Step 1

According to a previous study, the FINEs were distributed throughout the stratum basal, stratum spinosum, and terminated at the stratum granulosum [76], thus the sub-images of TFETM image stack from the layer of stratum granulosum (SG) to the dermis (D) were analyzed. The layer of stratum corneum (SC) was a thick layer of dead skin cells and the layer of stratum lucidum (SL) was a thin layer of dead skin cells that can only be seen on the palms and soles of the feet, with no FINEs distributed in the SC and the SL layers [77]. The definition of the skin layer of the TFETM image is as described in a previous study [78].

#### Step 2

In this step, each single-layer image from the SG to D layers was considered and analyzed. We briefly distinguished that FINEs signals appeared with dot-like and string-like patterns in human skin. Since the axon diameter of unmyelinated FINEs was morphologically observed from 0.2 to 1.5 µm based on electron micrograms (Introduction), thus the size definition of a dot-like pattern of the THG signal should be greater than 200 nm. On the other hand, the definition of a sting-like pattern was the THG signal consisting of dense dot-like patterns extended in one or multiple orientations. If the THG signal of a single-layer image is defined as a sting-like pattern, it could directly go into Step 4 of the operational protocol.

#### Step 3

Based on our IHC staining, the interval distance between each varicosity structure of unmyelinated FINEs was 0 to 10.4 μm (Results, Skin innervation of lower distal extremities tissue). In this step, we considered the situation in different layer images of the TFETM image stack. If the THG signal with a dot-like pattern extended along the same direction or a plurality of directions in different layer images, which means the signal was consecutive or the interval distance between each dot-like signal should be less than 10.4 μm. However, if the THG signal with a dot-like pattern didn’t extend or with an interval distance greater than 10.4 μm in different layer images, it cannot be defined as a FINEs signal.

#### Step 4

There are three conditions considered in this step. First, considering the circular contour of the cells, the signal arranged as a circle resembling cell signals in the skin was excluded. Second, to avoid mistaking FINEs for melanocyte dendrites, we excluded dendritic-cell-like THG signals. Third, the THG trace with less than 5 consecutive dot-like signals will not be considered. Consecutive signals are defined with a distance less than 10.4μm (Fig. 7 (i-k)).

#### Step 5

As shown in the E and F cases of Fig. 7 (h), if one fragment of FINEs signals is distributed in the upper epidermis (in stratum spinosum and stratum granulosum) and the other one distributed near the epidermal-dermal junction (EDJ), it is possible that these two fragments may belong to the same nerve fiber. If only one fragment of FINEs is located in the upper epidermis, it is possible that the nerves close to the EDJ layer may be distributed outside the stack due to the limited field of view (FOV). To avoid double counting, we considered the distance between the FINEs signal in the epidermis and the EDJ. When the closest distance between FINEs signals and EDJ is less than 5 μm, the FINEs structure inside the epidermis will be counted as one IENF.

### Quantification analysis of IENF index

The IENF index represents the IENF area density. The definition of IENF index is different from the IENF density of skin biopsy, which is a one-dimensional density. The IENF index of TFETM image is defined as:

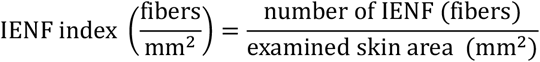

The examined skin area is equivalent to the total FOV of the TFETM images. Each IENF of the TFETM image with branching points in the epidermis is considered as one unit, while each IENF with branching points in the dermis is counted separately. The IENF index was derived and expressed as fibers/mm^2^. To avoid the deviation of nerve density statistics caused by the small acquisition area, the total FOV area of the TFETM image must be >1 mm^2^ for each subject. Moreover, we conducted two inspection procedures to ensure that all the *in vivo* TFETM image stacks with any longitudinal and lateral motion artifacts in the imaged epidermis were excluded as invalid. A preliminary inspection was performed immediately after the acquisition of the TFETM images. If an obvious motion blur was detected, the image stack was excluded, and the TFETM images were re-acquired. A second inspection was performed when analyzing the clinical data, and the TFETM images with lateral and/or axial motion artifacts in the epidermis were recorded and excluded.

### Calibration of IENF index

Given the different conditions of human skin in each volunteer, thus calibration factor that may affect the THG intensity of the TFETM image should be considered and calibrated for the IENF index. Lower THG intensities at the EDJ layer caused by the greater epidermis thickness also result in lower signal-background ratio (SBR) of FINEs signals. Due to the lack of other data and information about the relationship between nerves and the calibration factor, we made the simplest assumption that the relationship between calibration factor and the IENF index should be linear. Then, the IENF index was linearly calibrated according to this factor.

### Statistical analysis

All collected data were decoded and analyzed by blinded evaluators using the SPSS software (version 20.0; IBM Corporation, Armonk, NY, USA). The student’s t-tests were two-sided. *P*-values of <0.05 denoted statistically significant differences. Also, the Pearson correlation was performed to measure the strength of the linear relationship between two variables (IENF density & IENF index) in this study.

### Reporting summary

Further information on research design is available in the Nature Research Reporting Summary linked to this article.

## Data availability

For small animal experiment, the data that support the findings of this study are available on reasonable request from the corresponding author, C.K.Sun. For clinical trial, all the data are not redistributable to researchers other than those engaged in the IRB approved research collaborations from the author upon reasonable request. The data are not publicly available yet because that it contains information that could compromise privacy/consent. The corresponding author (W.Z. Sun) will however be able to consider specific requests on a case-by-case basis.

## Code availability

The ImageJ software (http://rsb.info.nih.gov/ij/) and the SPSS software (version 20.0; IBM Corporation, Armonk, NY, USA) was used for image/data processing. ImageJ software (http://rsb.info.nih.gov/ij/), whose source code is freely available, was written by Wayne Rasband at the National Institute of Health (Bethesda, MD, USA).

## Acknowledgment

We are indebted to the patients who participated in the clinical trials. This work was sponsored by National Science and Technology Council under MOST 107-2321-B-002-006, MOST 110-2221-E-002-048-MY3, and Ministry of Economic Affairs under 111-EC-17-A-19-S6-009.

## Author contributions

C.K.S., P.J.W., S.T.H, W.Z.S., and C.T. Y designed the experiment. P.J.W and H.C.T. built the TFETM system, performed the experiments, and analyzed the data. C.K.S. supervised the project. C.T.Y. and

W.Y.L. assisted the small animal experiment. S.T.H., C.C.C., and Y.H.L. recruited the volunteers. W.Z.S., S.T.H., C.C.C., and Y.H.L. conducted the clinical trial. S.T.H. and C.C.C. performed the skin biopsy. All authors were involved in discussions during the work and in the preparation of the manuscript.

## Additional information

*Funding.* National Science and Technology Council, Taiwan (MOST 107-2321-B-002-006, MOST 110-2221-E-002-048-MY3) & Ministry of Economic Affairs (111-EC-17-A-19-S6-009).

